# Increased *H. pylori* stool shedding and EPIYA-D *cagA* alleles are associated with gastric cancer in an East Asian hospital

**DOI:** 10.1101/284448

**Authors:** Sarah Talarico, Christina Leverich, Bing Wei, Jie Ma, XinGuang Cao, YongJun Guo, GuangSen Han, Lena Yao, Steve Self, Yuzhou Zhao, Nina R. Salama

## Abstract

**Background:** *Helicobacter pylori* infection induces chronic inflammation and tissue damage in the stomach, increasing risk for gastric cancer. Paradoxically, these tissue alterations may promote loss of *H. pylori* infection during cancer progression. *H. pylori*’s role in cancer progression beyond initiation is unclear. Geographic variation in gastric cancer risk has been attributed to variation in carriage and type of the *H. pylori* oncogene *cagA.*

**Methods:** To investigate possible differences in *H. pylori* load in the stomach and shedding in stool, *H. pylori* load and *cagA* genotype were assessed using droplet digital PCR assays on gastric mucosa and stool samples from 49 urea breath test-positive individuals, including 25 gastric cancer and 24 non-cancer subjects at Henan Cancer Hospital, Henan, China.

**Results:** Quantitation of *H. pylori* DNA indicated similar gastric loads among cancer and non-cancer cases, but the gastric cancer group had a median *H. pylori* load in the stool that was six times higher than that of the non-cancer subjects. While the *cagA* gene was uniformly present among study subjects, only 70% had the East Asian *cagA* allele, which was significantly associated with gastric cancer (Fisher’s Exact Test, p = 0.03).

**Conclusion:** *H. pylori* persists in a subset of gastric cancer cases and thus may contribute to cancer progression. In this East Asian population with a high prevalence of the *cagA* gene, the East Asian allele could still provide a marker for gastric cancer risk.

**Impact:** This study contributes to our understanding of *H. pylori* dynamics in the context of pathological changes.

## Introduction

The bacterial pathogen *Helicobacter pylori* infects the human stomach causing chronic inflammation of the gastric mucosa. Most infected individuals remain asymptomatic but a subset develop peptic ulcers, gastric adenocarcinomas, and mucosa-associated lymphoid tissue (MALT) lymphoma resulting from the infection. Disease risk depends on the severity and distribution of inflammation in the stomach. Those with inflammation that is predominantly in the antrum of the stomach are at higher risk for duodenal ulcer and those with inflammation that extends into the stomach corpus are at higher risk for gastric adenocarcinoma (1). In those who eventually develop gastric adenocarcinoma, *H. pylori*-induced chronic inflammation initiates a pathological cascade that progresses from atrophy to intestinal metaplasia and dysplasia (2). Density of *H. pylori* infection has been found to be correlated with level of inflammation (3, 4) but loss of *H. pylori* infection has been observed in atrophic gastric tissue (5) and increased pH in the stomach can promote growth of other bacteria (6). Notably, *H. pylori* often cannot be detected within tumors and the association of *H. pylori* with cancer was strongest when *H. pylori* infection was assessed 10 years prior to cancer diagnosis (7).

Bacterial genetic factors contribute in part to differences in bacterial density, inflammation, and disease development. The strain-variable *cagA* gene is associated with a higher risk of gastric adenocarcinoma development (8). The *cagA* gene is located within the *cag* pathogenicity island that encodes a Type IV secretion system which delivers the CagA effector protein into host gastric epithelial cells (9). Inside the host cell, the CagA protein is phosphorylated at EPIYA sites and the phosphorylated form is able to deregulate normal cellular signaling (10-13). The *cagA* gene is grouped into two different allele types, Western alleles that encode an EPIYA-C motif and East Asian alleles that encode an EPIYA-D motif. *cagA* alleles having an EPIYA-D motif are predominantly found in *H. pylori* strains circulating in East Asian countries and are associated with an increased risk of gastric cancer development compared to *cagA* alleles having an EPIYA-C motif (14).

Studies investigating the role of the *cagA* gene in density of *H. pylori* infection in the stomach have produced conflicting results, with some studies reporting a significantly higher density of *H. pylori* infection in those with *cagA*-positive *H. pylori* (3, 15) and some finding no difference in bacterial density between *cagA*-positive and *cagA*-negative strains (4, 16, 17). We recently reported development of a new, non-invasive method for detection, quantification, and *cagA* genotyping of *H. pylori* from stool samples that uses droplet digital PCR (ddPCR). We tested this method using a collection of matched serum and stool samples from volunteers in Costa Rica and observed a two log range of *H. pylori* loads in the stool. Furthermore, the *H. pylori* load in the stool was significantly higher in those with a *cagA*-positive strain (18). However, the extent to which the *H. pylori* load in the stool reflects the load in the stomach is not clear.

To assess differences in *H. pylori* carriage and colonization density upon cancer development and to examine factors that contribute to the observed variability in *H. pylori* loads in stool, including *H. pylori* load in the stomach and *cagA* status, we applied the quantitative ddPCR assays to gastric mucosal brushing samples and stool samples collected from gastric cancer and non-cancer subjects from China.

## MATERIALS AND METHODS

### Study Populations and Specimen Collection

To investigate possible differences in *H. pylori* carriage and load in the stomach as well as *H. pylori* shedding in stool, C14 urea breath test (UBT, Haidewei HUBT-01P) was used to screen individuals to be treated for gastric cancer or undergoing upper GI endoscopy for other indications at Henan Cancer Hospital between 2015.10.27-2016.03.15. Subjects who were *H. pylori*-positive by UBT were offered participation in this research study and all participants provided written informed consent prior to participation. Participating subjects had gastric mucosal brushings collected and were asked to fill out a questionnaire that covered demographic characteristics, medical conditions, and medications and provide a stool sample. All procedures were approved by the Henan Cancer Hospital Medical Research Institution Review Board (doc # 2016oct005).

In total, samples were collected from 50 subjects including 25 gastric cancer subjects diagnosed by pathology who were undergoing gastric resection surgery and 25 non-cancer subjects with no history of gastric tumor or surgery who were undergoing upper GI endoscopy either as indicated because of symptoms or as asymptomatic volunteers. All non-cancer cases showed histologic evidence of gastritis and no ulcers. All 50 subjects had not taken antibiotics for gastric disease in the past month.

Gastric mucosal brushing samples were collected from two different anatomical sites in the stomach, the antrum and the corpus, using cytology brushes (Puritan Medical Products Co LLC). Gastric mucosal brushings were collected and stored in a cryovial containing 1 ml flow media (minimal essential media plus 10% DMSO, 5% fetal calf serum, 5mM HEPES (19). Cryovials containing the gastric mucosal brushing samples were immediately placed and kept at −80°C. The gastric mucosal brushings were collected from the gastric cancer subjects during the gastric resection surgery and from non-cancer subjects during upper GI endoscopy. Stool samples were collected 1-2 days before either surgery or endoscopy. Participants undergoing surgery collected the stool sample at the hospital and participants undergoing endoscopy collected the stool sample at home and delivered it to the hospital on the same day. Stool samples were collected by the participants into a vial containing 5 ml RNAlater nucleic acid preservative (Ambion) and were frozen at −20°C upon receipt. In addition to the gastric mucosal brushings and stool samples, gastric biopsies were collected for histological analysis. A gastric biopsy was collected from the antrum and the corpus. Gastric biopsies were formalin-fixed and paraffin-embedded prior to histological analysis.

### DNA Extraction

For the gastric mucosal brushings, sample media and brush were transferred from the cryovial to a microcentrifuge tube and centrifuged, followed by removal of the supernatant and brush. DNA was then extracted from the pellet using the UltraClean Tissue and Cells DNA Isolation Kit (MoBio) according to the manufacturer’s instructions.

Stool samples were first transferred to a microcentrifuge tube and centrifuged to remove the RNAlater. DNA was then extracted using the QIAamp Stool DNA Mini Kit (Qiagen) according to the manufacturer’s instructions, with the lysis step performed at 95°C. Stool DNA concentration was measured using a NanoDrop 2000 UV-Vis Spectrophotometer (Thermo Scientific), and the concentration was adjusted to 100 ng/µl.

### Droplet Digital PCR

*H. pylori*-specific droplet digital PCR (ddPCR) assays (*H. pylori* 16S assay, *cagA* detection assay, and *cagA* EPIYA typing assay) were performed using the QX200 ddPCR System (BioRad) as described previously (18) for stool DNA and gastric mucosal brushing DNA. Briefly, each 20 µl reaction contained 1x ddPCR Supermix for Probes (BioRad), 900 nM of each primer, 250 nM of probes, and 10 µl DNA. Reactions were subject to thermal cycling with the following conditions: 95°C for 10 minutes, 45 cycles of 94°C for 30 seconds and 55°C for 1 minute, and a final incubation at 98°C for 10 minutes. Data were analyzed using the QuantaSoft software version 1.6.6 (BioRad). The threshold was set to 4500 for the *H. pylori* 16S assay and 5500 for the *cagA* detection assay. For the *cagA* EPIYA assay, the thresholds were set to 4000 (both gastric and stool samples) for the FAM channel and 2500 (gastric samples) or 3000 (stool samples) for the HEX channel. A positive control (stool DNA from an *H. pylori*-positive volunteer) and negative control (molecular grade water) were included in each batch of samples analyzed.

For the gastric brushing samples, the copy number of *H. pylori* 16S was normalized to the copies of human DNA in the sample, as measured by a ddPCR assay for 18S. The following primers and probe were used in the 18S assay: 18SFor (5’ -CGATGCTCTTAGCTGAGTG-3’), 18SRev (5’ -CTTAATCATGGCCTCAGTTC -3’), and 18S_HEX (5’ hexachloro-fluorescein -CCGCAGCTAGGAATAATGGAATAG – black hole quencher 3’). Reactions were subject to thermal cycling with the following conditions: 95°C for 10 minutes, 45 cycles of 94°C for 30 seconds and 59°C for 1 minute, and a final incubation at 98°C for 10 minutes. Data were analyzed using the QuantaSoft software version 1.6.6 (BioRad) and the threshold was set to 2000.

The concentration of the gastric mucosal brushing DNA sample was adjusted so that the copy number of the ddPCR assay target was within the dynamic range of the assay. DNA samples were run as either undiluted or diluted 1:10, 1:100, 1:1000, or 1:10,000. Stool DNA concentration was adjusted to 100 ng/µl as needed so that no more than 1 µg stool DNA was added to each ddPCR reaction.

### Histological Analysis

For histological analysis of the gastric biopsies, slides were made from formalin-fixed paraffin-embedded (FFPE) gastric tissue samples. Slides were stained with Wright-Giemsa stain (Baso Diagnostics, Inc. Zhuhai, China) according to the manufacturer’s instructions. Stained slides were analyzed and scored as either normal (0), mild (1), moderate (2), or marked (3) according to the Sydney Classification System for *H. pylori* density, neutrophils, mononuclear cells, atrophy, and intestinal metaplasia (20). For the non-cancer subjects, more than one anatomical site of the stomach was analyzed for a patient (i.e. antrum and corpus) and the *H. pylori* density, neutrophils, mononuclear cells, and intestinal metaplasia was scored as the average of the two sites. Atrophy was scored separately for the antrum and corpus. For the gastric cancer subjects, only gastric tissue from the carcinoma adjacent site was scored.

### Statistical Analysis

Correlation between gastric cancer and *cagA* allele was assessed using Fisher’s Exact Test. The Wilcoxon Two-Sample Exact Test was used for comparisons of *H. pylori* loads between gastric cancer subjects and non-cancer subjects and between East Asian *cagA* allele and Western *cagA* allele groups. Correlation between *H. pylori* load in the gastric mucosal brushing samples from different anatomic sites and with *H. pylori* load in the stool was analyzed using Spearman Correlation Coefficients with Fisher’s z-Transformation. All statistical analyses were performed in SAS version 9.4 (SAS Institute, Inc.).

## RESULTS

### Gastric Cancer cases show frequent evidence of active H. pylori infection

Of 58 gastric cancer subjects screened for *H. pylori* infection by UBT, 25 (43%) were *H. pylori*-positive. Of 108 non-cancer subjects screened, 25 (23%) were *H. pylori*-positive by UBT. One of the 25 participating non-cancer subjects did not provide an adequate stool sample and was excluded from the analysis. The remaining 49 subjects included in our study had a median age of 53 years (range 27 to 76 years) and included 15 (31%) females and 34 (69%) males. Of the 25 gastric cancer subjects, 22 (88%) were male compared to 12 (50%) of the 24 non-cancer cases. The gastric cancer cases had a median age of 59 years (range 46 – 76 years) and the non-cancer cases had a median age of 49 years (range 27 – 66 years).

Of the 49 subjects, histological slides of gastric biopsy were available for all 25 of the gastric cancer subjects and 17 of the non-cancer subjects consented to provide samples for histologic scoring of inflammation, pathological changes, and *H. pylori* density by the Sydney Classification System (20). Level of neutrophil and mononuclear cell infiltration, indicating acute and chronic inflammation respectively, tended to be lower in the gastric cancer subjects, but distribution of scores for atrophy, intestinal metaplasia, and *H. pylori* density were similar between gastric cancer subjects and non-cancer subjects (Table 1). Of the 42 subjects with histology slides analyzed, 13 (31%) had signs of atrophy and 18 (43%) had signs of intestinal metaplasia. Despite all subjects being positive for *H. pylori* infection by UBT, three subjects (two cancer cases and one non-cancer subject) did not have *H. pylori* detected by histology.

**Table 1.** Sydney Classification System scores for gastric histology slides from 25 gastric cancer subjects and 17 non-cancer subjects from Henan Cancer Hospital, China

|  | Score<sup>a</sup> | No. (%)<br>Gastric<br>Cancer<br>Subjects<br>(n = 25) | No. (%)<br>Non-cancer<br>Subjects<br>(n = 17) |
| --- | --- | --- | --- |
| Neutrophils | 0 | 10 (40) | 2 (12) |
|  | 1 | 9 (36) | 11 (65) |
|  | 2 | 6 (24) | 4 (24) |
|  | 3 | 0 | 0 |
| Mononuclear Cells | 0 | 0 | 0 |
|  | 1 | 16 (64) | 4 (24) |
|  | 2 | 7 (28) | 5 (29) |
|  | 3 | 2 (8) | 8 (47) |
| Atrophy | 0 | 17 (68) | 12 (71) |
|  | 1 | 7 (28) | 3 (18) |
|  | 2 | 1 (4) | 2 (12) |
|  | 3 | 0 | 0 |
| Intestinal Metaplasia | 0 | 13 (52) | 11 (65) |
|  | 1 | 8 (32) | 5 (29) |
|  | 2 | 3 (12) | 1 (6) |
|  | 3 | 1 (4) | 0 |
| *H. pylori* | 0 | 2 (8) | 1 (6) |
|  | 1 | 13 (52) | 7 (41) |
|  | 2 | 6 (24) | 6 (35) |
|  | 3 | 4 (16) | 3 (18) |

<sup>a</sup> 0 = normal, 1 = mild, 2 = moderate, 3 = marked; score is for the carcinoma adjacent tissue of gastric cancer subjects and is the average of the antrum and corpus for non-cancer subjects

### *Cancer and non-cancer cases show similar loads of* H. pylori *in the stomach by ddPCR*

To investigate the load of *H. pylori* in the stomach among subjects, we employed ddPCR to measure the copy number of the *H. pylori* 16S ribosomal RNA gene, a highly conserved gene present in all *H. pylori* strains, in total DNA extracted from mucosal brush samples from both the antrum and corpus for each subject. *H. pylori* load was normalized to the human 18S ribosomal RNA gene copy number present in each sample. The *H. pylori* 16S gene was detected in one or both gastric mucosal brushing samples of 22 (88%) of 25 gastric cancer subjects and all 24 (100%) non-cancer subjects. One non-cancer subject had *H. pylori* 16S detected in the sample from the corpus but not the antrum. Of the three subjects that did not have *H. pylori* detected by histology, one also did not have the *H. pylori* 16S gene detected by ddPCR in either gastric sample. The other two had low loads of *H. pylori* 16S detected by ddPCR (gastric *H. pylori* load of 10 and 0.4 16S copies per 1000 18S). As shown in Figure 1A, the load of *H. pylori* detected in gastric samples varied over several logs in both cancer and non-cancer subjects. For the gastric samples with detectable *H. pylori* 16S, the *H. pylori* 16S copy number per 1000 18S ranged from 0.003 to 1034 with a median of 144 for the antrum and ranged from 0.84 to 1967 with a median of 103 for the corpus. The *H. pylori* load in the antrum and corpus were significantly correlated (Spearman Correlation Estimate = 0.7, 95% CI: 0.5 – 0.8, Figure 1B) and for subsequent analysis we considered the gastric load to be the average of that measured for the antrum and corpus sample for each subject. The gastric *H. pylori* load between the gastric cancer cases (median=139 *H. pylori* 16S copies per 1000 18S copies, range 0.9 – 386) and the non-cancer subjects (median=114 *H. pylori* 16S copies per 1000 18S copies, range 0.4 −1150) was not significantly different (p=0.4, Wilcoxon Exact Test).

**Figure 1.**
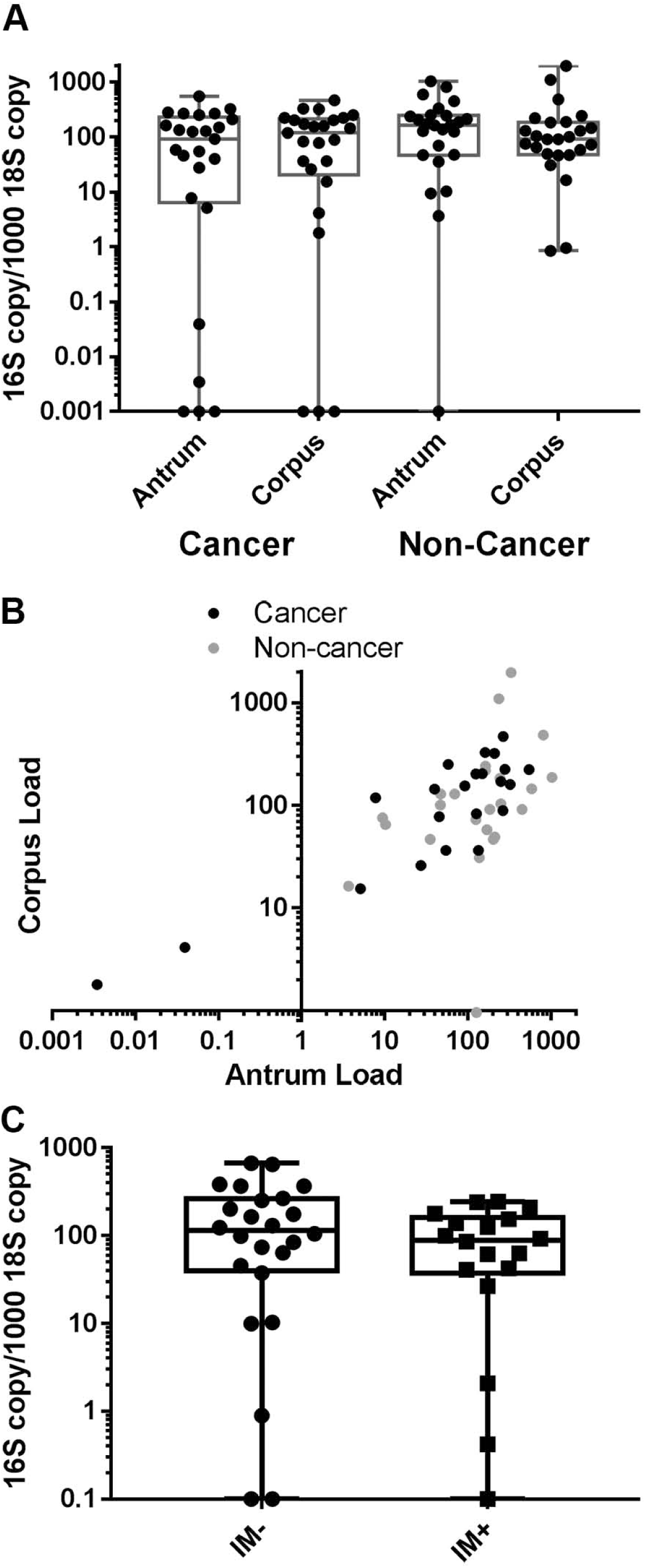
Gastric cancer cases and non-cancer subjects show similar variation in *H. pylori* load within the stomach and high concordance between antrum and corpus samples. **A.** Copy number of the *H. pylori* 16S ribosomal RNA genes was measured by ddPCR from gastric brushes collected from the stomach antrum or corpus and normalized to copy number of the human 18S ribosomal RNA gene. Cancer cases n=25, non-cancer subjects n=24. Box extends from 25^th^ to 75^th^ percentiles with a line at the median value. Whiskers show minimum and maximum values. In three cancer cases, no *H. pylori* DNA was detected and in one non-cancer subject, *H. pylori* was only detected in the corpus. These points are plotted on the X-axis. **B.** Correlation between corpus and antrum load (copy number of *H. pylori* 16S ribosomal gene) for subjects with *H. pylori* DNA detected in both samples. Cancer cases black, n=22, non-cancer subjects grey, n=23. Spearman Correlation Estimate = 0.7 (95% CI: 0.5 – 0.8). **C.** Gastric load (average of corpus and antrum) in subjects with (IM+, n=18) or without (IM-, n=24) histologic evidence of intestinal metaplasia (IM). Biopsy samples for histology were not available for 7 non-cancer cases. Points plotted on the X-axis represent subjects for which no *H. pylori* 16S rRNA gene was detected.

Loss of acid secreting parietal cells and transformation of the gastric glands to an intestinal metaplasia phenotype as consequence of *H. pylori* induced inflammation has been suggested to favor outgrowth of other bacterial species over *H. pylori*. We saw similar proportions of intestinal metaplasia in cancer cases compared to non-cancer subjects (Table1). We thus checked whether gastric *H. pylori* load differed according to presence of intestinal metaplasia (Fig. 1C), but saw no significant difference (p=0.3, Wilcoxon Exact Test).

### *Stool-based ddPCR detection of* H. pylori 16S *is sensitive, but shows little correlation with gastric load*

We next examined the relationship between detection and copy number of the *H. pylori* 16S ribosomal RNA gene by ddPCR in stool with detection and copy number in the stomach. Of the stool samples from gastric cancer cases, the *H. pylori* 16S gene was detected in 21 (84% of the 25 gastric cancer cases). Considering the 22 gastric cancer cases for which we detected the *H. pylori 16S* gene in the stomach, 21 (95%) also had detection in the stool. The *H. pylori 16S* gene was detected in 22 (92%) of the 24 stool samples from non-cancer subjects. For the stool samples with detectable *H. pylori 16S*, the *H. pylori 16S* copy number per µg stool DNA ranged from 2 to 1080 with a median of 26 (Figure 2A). The *H. pylori* load in the stool was significantly higher (Wilcoxon Exact Test, p= 0.03) in the gastric cancer cases (median=46 *H.pylori* 16S copies per µg stool DNA, range 0 -560) compared to the non-cancer subjects (median=7.5 *H.pylori* 16S copies per µg stool DNA, range 0 -1080). The *H. pylori* load observed in the stool was only weakly correlated with the *H. pylori* load in the gastric samples of the same subject (Spearman Correlation Estimate = 0.3, 95% CI:0.03 – 0.6, Figure 2B).

**Figure 2.**
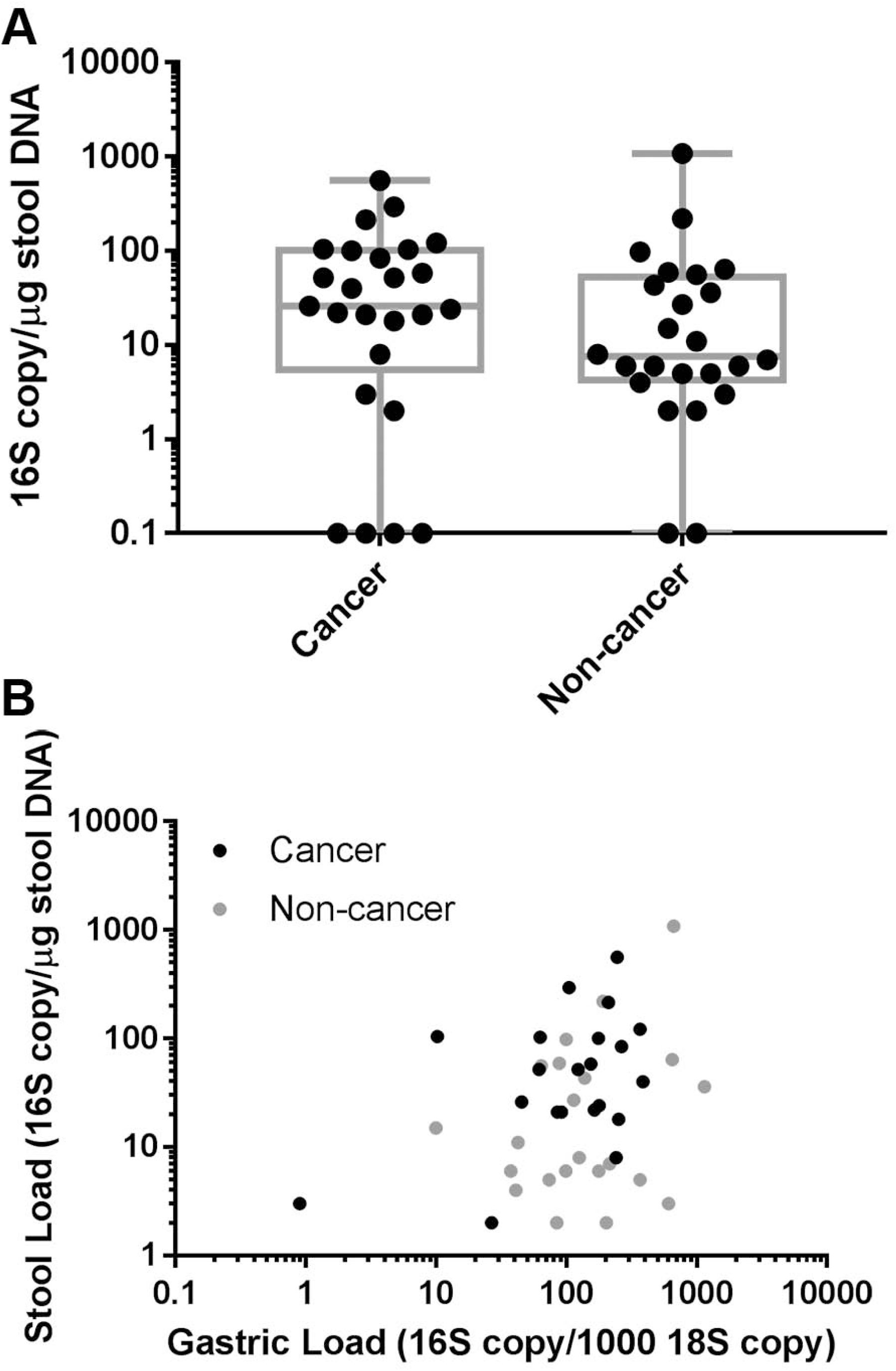
*H. pylori* DNA is detected in stool from both cancer and non-cancer subjects. **A.** Copy number of *H. pylori* 16S DNA was measured by ddPCR of total stool DNA and normalized to µg stool DNA. Box extends from 25^th^ to 75^th^ percentiles with a line at the median value. Whiskers show minimum and maximum values. Cancer cases n=25, non-cancer cases n=24. **B.** Copy number of *H. pylori* 16S DNA was measured by ddPCR and normalized to µg total stool DNA (stool load) or copy number of human 18S (gastric load). Gastric load calculated as the average of the corpus and antrum samples for each subject. Cancer cases n=22, non-cancer cases n=24. One cancer and two non-cancer subjects had no *H. pylori* DNA detected in stool and are not plotted. Spearman Correlation Estimate = 0.3, 95% CI: 0.03 – 0.6.

### *ddPCR reveals variation in* cagA *alleles and association of East Asian cagA with gastric cancer*

Previous studies suggest very high prevalence of *cagA*-positive strains and a predominance of East Asian EPIYA alleles in China (21). To assess the presence of the *cagA* gene and distinguish between East Asian and Western alleles, we performed our *cagA* detection and *cagA* EPIYA typing ddPCR assays (18) on the antrum and corpus samples. Of the 46 subjects with detectable *H. pylori* 16S, all 46 were positive for the *cagA* gene by one or both assays. Of these 46 subjects, the *cagA* EPIYA typing ddPCR classified 37 (80%) as having either an East Asian or Western *cagA* allele. PCR amplification and sequencing of a portion of the *cagA* gene was performed for seven antrum samples that were negative by the *cagA* EPIYA typing assay. Of these seven samples, five were determined by sequencing to have EPIYA-C (Western *cagA* allele) and two had neither EPIYA-D nor EPIYA-C. The five that were negative for the *cagA* EPIYA typing ddPCR assay but were determined to have EPIYA-C by sequencing all had the same three nucleotide differences in the forward primer region that could explain failure of the assay for these samples. One subject was determined to have an East Asian allele in the corpus sample by the *cagA* EPIYA typing ddPCR assay and an allele having neither an EPIYA-D nor an EPIYA-C in the antrum sample by sequencing.

Combining the results of the *cagA* EPIYA typing ddPCR assay and sequencing for the 46 subjects, 32 (70%) had the East Asian allele, 10 (22%) had a Western allele, one had only a non-East Asian, non-Western allele, and three were not determined. Being infected with an *H. pylori* strain with the East Asian *cagA* allele was significantly associated with gastric cancer (Fisher’s Exact Test, p = 0.03) (Table 2). Since we observed higher *H. pylori* loads in the stool among gastric cancer cases, we examined the relationship between *H. pylori* load in the stool and *cagA* allele. The median *H. pylori* stool load for subjects with an East Asian *cagA* allele was 33.5 *H. pylori* 16S copies per µg stool DNA (range 0 – 560) and for subjects with a Western *cagA* allele was 15.5 *H. pylori* 16S copies per µg stool DNA (range 0 – 1080) (Wilcoxon Exact Test p = 0.26).

**Table 2.**
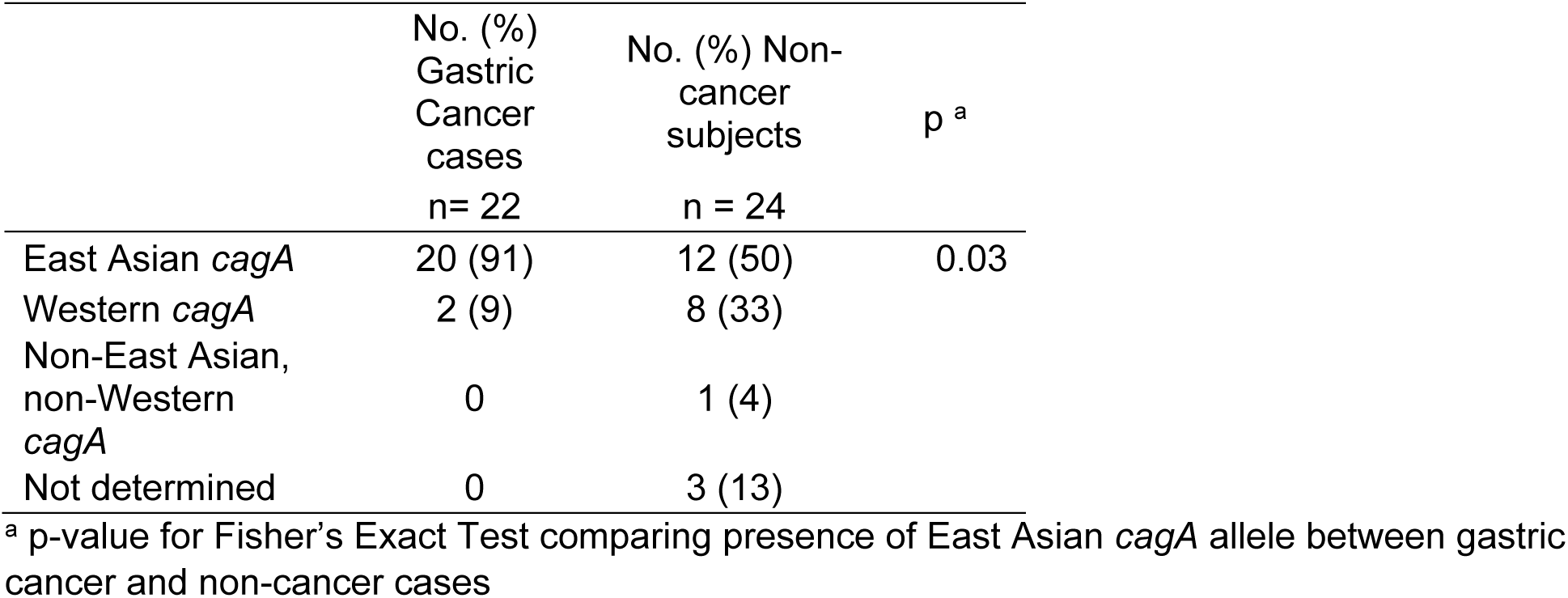
Comparison of *cagA* genotype between gastric cancer cases and non-cancer subjects.

|  | No. (%)<br>Gastric<br>Cancer<br>cases<br>n= 22 | No. (%) Non-cancer<br>subjects<br>n = 24 | p<sup>a</sup> |
| --- | --- | --- | --- |
| East Asian *cagA* | 20 (91) | 12 (50) | 0.03 |
| Western *cagA* | 2 (9) | 8 (33) |  |
| Non-East Asian,<br>non-Western<br>*cagA* | 0 | 1 (4) |  |
| Not determined | 0 | 3 (13) |  |

<sup>a</sup> p-value for Fisher's Exact Test comparing presence of East Asian *cagA* allele between gastric cancer and non-cancer cases

### *ddPCR-based* cagA-*detection in stool samples shows high specificity but lower sensitivity compared to stomach samples*

We extended our analysis of *cagA* genotypes by running the *cagA* EPIYA ddPCR typing assay on stool samples from 48 subjects. For one endoscopy case, there was insufficient stool DNA to run our *cagA* EPIYA ddPCR assay. We detected *cagA* in stool for 23/31 (73%) of subjects where we detected EPIYA-D type *cagA* in the stomach and 4/5 (80%) of subjects where we detected EPIYA-C type *cagA* in the stomach by the *cagA* EPIYA ddPCR assay. In all cases we detected the same EPIYA type in the stomach and stool samples by the *cagA* EPIYA ddPCR assay.

## DISCUSSION

In this study, we leveraged recently developed ddPCR-based quantitative assays to explore differences in *H. pylori* load present in the stomach and shed into stool among cancer and non-cancer cases undergoing surgery or upper GI endoscopy respectively at a hospital in China’s Henan province, a region with high incidence of stomach cancer (38.13 per 100,000). At present, all methods for detecting active *H. pylori* infection (UBT, histology, PCR, culture) have limitations in specificity and sensitivity such that many studies require concordance between two or more methods. We used the clinically accepted UBT to screen for cases with *H. pylori* infection that were then analyzed by histology and ddPCR. Using UBT, our observations of 43% and 23% prevalence of *H. pylori* infection in cancer and non-cancer cases falls at the low end of that observed among symptomatic individuals in similar regions of China (30-80%) (22-24). Of the UBT-positive cases enrolled in our study, 89% were positive by all three assays. While tissue alterations associated with cancer progression have been suggested to disfavor *H. pylori* colonization, we found similar loads in the stomach among cancer cases and non-cancer cases. Additionally, histologic scores were not significantly different among cancer cases and non-cancer cases. The high correlation between loads measured from different biopsy samples and regions of the stomach within individuals suggests that patchy colonization within the stomach may be the exception rather than the rule in this population. Furthermore, the continued presence of *H. pylori* at high levels in the stomach may influence the rate or molecular phenotype of gastric cancer in the subset of individuals that remain colonized.

In contrast to the similar loads observed in the stomach, gastric cancer cases showed higher *H. pylori* loads in stool. Our previous population-based study in Costa Rica measured higher *H. pylori* loads in stool among subjects positive for CagA by serology (18), prompting us to assay *cagA* status. However, all 46 subjects that had *H. pylori* 16S detected in the gastric samples also had *cagA* detected, so it was not possible to analyze the correlation between presence of *cagA* and *H. pylori* load in the stool in this population. While the East Asian *cagA* allele was significantly associated with gastric cancer, the allele was not associated with a higher *H. pylori* load in the stool. This suggests that the higher *H. pylori* load in subjects with gastric cancer is not dependent on presence of the *cagA* gene or East Asian *cagA* allele. Further studies will be necessary to determine at what point during progression to gastric cancer infected individuals have a higher *H. pylori* load in the stool and whether this could be used as a marker either for presence of gastric cancer or risk for eventual gastric cancer development.

In this study in Henan, China, we found the *cagA* gene to be uniformly present among study subjects but only 70% had the East Asian *cagA* allele, which was significantly associated with gastric cancer. Previous studies examining the *cagA* gene and East Asian allele in Chinese populations have found the prevalence of the *cagA* gene to be very high (75 – 90%) with the East Asian allele being predominant (25, 26). While the *cagA* gene has been a good marker in Western populations for cancer risk, its utility in East Asian populations has been less clear since most strains have the *cagA* gene. In this population with a high prevalence of the *cagA* virulence gene, the East Asian allele could still provide a marker for gastric cancer risk.

The limited correlation between *H. pylori* loads measured in the stomach compared to the stool suggests shedding from the stomach depends on more than absolute abundance in the stomach. Possible mechanisms include altered growth rate, induction of epithelial turnover, and survival during transit through the lower GI tract. Further studies will need to be done to investigate the factors that contribute to the observed higher *H. pylori* load in the stool among gastric cancer subjects compared to non-cancer subjects despite a similar *H. pylori* load in the stomach. This will further our understanding of how pathological changes associated with gastric cancer development affect *H. pylori* colonization as well as what continued contribution *H. pylori* has to gastric cancer beyond initiation.

